# clustifyr: An R package for automated single-cell RNA sequencing cluster classification

**DOI:** 10.1101/855064

**Authors:** Rui Fu, Austin E. Gillen, Ryan M. Sheridan, Chengzhe Tian, Michelle Daya, Yue Hao, Jay R. Hesselberth, Kent A. Riemondy

## Abstract

**Background:** In single-cell RNA sequencing (scRNA-seq) analysis, assignment of likely cell types remains a time-consuming, error-prone, and biased process. Current packages for identity assignment use limited types of reference data, and often have rigid data structure requirements. As such, a more flexible tool, capable of handling multiple types of reference data and data structures, would be beneficial.

**Findings:** To address difficulties in cluster identity assignment, we developed the clustifyr R package. The package leverages external datasets, including gene expression profiles from scRNA-seq, bulk RNA-seq, microarray expression data, and/or signature gene lists, to assign likely cell types. We benchmark various parameters of a correlation-based approach, and also implement a variety of gene list enrichment methods. By providing tools for exploratory data analysis, we demonstrate the feasibility of a simple and effective data-driven approach for cell type assignment in scRNA-seq cell clusters.

**Conclusions:** clustifyr is a lightweight and effective cell type assignment tool developed for compatibility with various scRNA-seq analysis workflows. clustifyr is publicly available at https://github.com/rnabioco/clustifyr

## INTRODUCTION

Single-cell mRNA sequencing promises to deliver improved understanding of cellular mechanisms, cell heterogeneity within tissue, and developmental transitions[1–5]. A key challenge in scRNA-seq data analysis is the identification of cell types from single-cell transcriptomes. Manual inspection of the expression patterns from a small number of marker genes is still standard practice, which is cumbersome and frequently inaccurate. Unfortunately, current implementations of scRNA-seq suffer from several limitations[3,6,7] that further compound the problem of cell type identification. One, only RNA levels are measured, which may not correlate with cell surface marker or gene expression signatures identified through other experimental techniques. Two, due to the low capture rate of RNAs, low expressing genes may face detection problems regardless of sequencing depth. Many previously established markers of disease or developmental processes suffer from this issue, such as transcription factors. On the data analysis front, over or under-clustering may generate cluster markers that are uninformative for cell type labeling. In addition, cluster markers that are unrecognizable to an investigator may indicate potentially interesting unexpected cell types, but can be very intimidating to interpret.

For these reasons, many investigators struggle to integrate scRNA-seq into their studies due to the challenges of confidently identifying previously characterized or novel cell populations.

Formalized data-driven approaches for assigning cell type labels to clusters will greatly aid researchers in interrogating scRNA-seq experiments. Currently, multiple cell type assignment packages exist but they are specifically tailored towards input types or workflows[8–10].

We developed the R package clustifyr, a lightweight and flexible tool that leverages a wide range of prior knowledge of cell types to pinpoint target cells of interest or assign general cell identities to difficult-to-annotate clusters. Here, we demonstrate its applications with transcriptomic information of external datasets and/or signature gene profiles, to explore and quantify likely cell types. The clustifyr package is built with compatibility and ease-of-use in mind to support other popular scRNA-seq tools and formats.

## METHODS

### Extracting information from existing R objects

For clustifyr, query data and reference data can take the form of raw or normalized expression matrices and corresponding metadata tables. To better integrate with standard workflows that involve S3/S4 R objects, methods for clustifyr are written to directly recognize Seurat[11] or SingleCellExperiment[12] objects, retrieve the required information, and reinsert classification results back into an output object (Fig. 1A). A more general wrapper is also included for compatibility with other common data structures, and can be easily extended to new object types.

**FIGURE 1.**
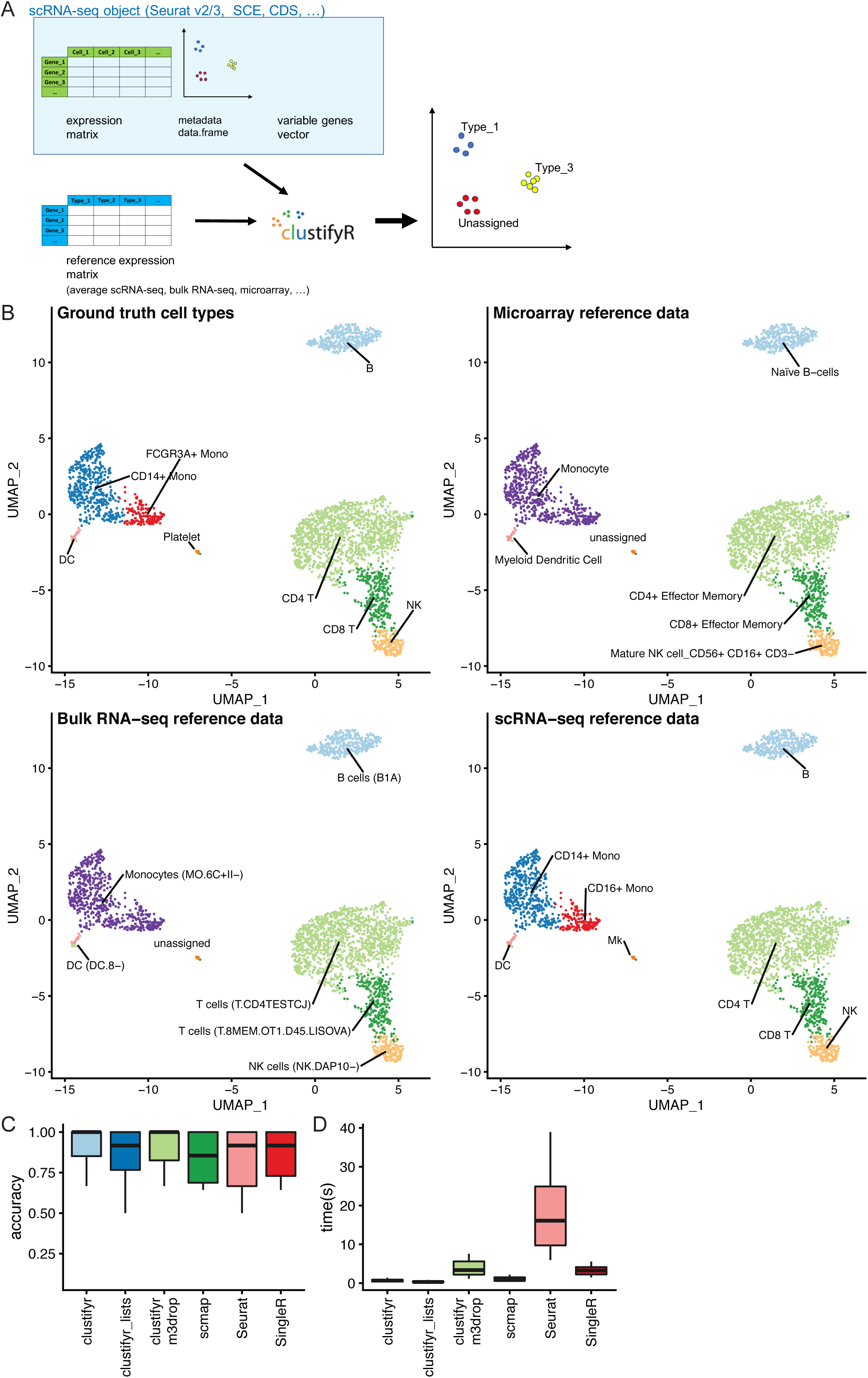
clustifyr uses many types of expression data for cluster identity assignment. A) Schematic of input data types supported by clustifyr. B) UMAP projections of PBMC cells colored by known cell type (Ground truth cell types) or cell types assigned by clustifyr using reference transcriptome data from microarray, sorted bulk RNA-seq, and scRNA-seq experiments. C) Accuracy of classifications generated by clustifyr or existing methods using the Tabula Muris to benchmark cell type classifications across sequencing platforms. D) Run-time of clustifyr or existing methods on the Tabula Muris cross-platform classification.

This approach also has the added benefit of forgoing certain calculations such as variable gene selection or clustering, which may already be stored within input objects. clustifyr is designed to perform per-cluster or per-cell classification after previous steps of analysis by other informatics tools. Therefore, it relies on, and is agnostic to, common external packages for cell clustering and variable feature selection. It has been tested against scRNA-seq data analyzed by Seurat[11] and Bioconductor SingleCellExperiment (SCE)[12]. We envision it to be compatible with all scRNA-seq processing, clustering, and marker gene discovery workflows. Simple and non-package-dependent functions for k-means clustering and selection of high variance genes are implemented as alternatives.

### Measuring correlation and comparing gene lists

To assess similarity between query and reference cell types, Spearman, Pearson, Kendall, and Cosine correlation calculations are implemented in clustifyr. Multiple methods are implemented to assess cell identity based on curated gene lists including hypergeometric tests, Jaccard Index, GSEA via the fgsea R package[13], mean percentage of cells that express marker genes, and marker scoring based on mean per-cell Spearman ranked correlation.

### Benchmarking

clustifyr was tested against scmap v1.8.0[8], SingleR v1.0.1[9], and Seurat v3.1.1[11]. scRNA-seq Tabula Muris data was downloaded from https://tabula-muris.ds.czbiohub.org/ as seuratV2 objects. Human pancreas data was downloaded from https://hemberg-lab.github.io/scRNA.seq.datasets/ as SCE objects. In all instances, to mimic the usage case of clustifyr, clustering and dimension reduction projections are acquired from available metadata, in lieu of new analysis.

An R script was modified to benchmark clustifyr following the approach and data sets of scRNAseq_Benchmark[14], using M3Drop[15] variable gene selection for every test. R code used for benchmarking, and preprocessing of other datasets, in the form of matrices and tables, are documented in R scripts available in the clustifyr GitHub repository.

## FINDINGS

Prior knowledge of cells types should facilitate cell identity assignment in scRNA-seq analysis. However, in practice, differences between flow cytometry, microarray data, bulk RNA-seq, and the implementations of scRNA-seq, including but not limited to Dropseq, Microwell-seq, 10X genomics 3’ end seq, and 5’ end seq, make cross-platform comparisons difficult. We therefore set out to build a flexible framework that could compare single-cell transcriptomes across different experimental methods.

Using clustifyr, which adopts correlation-based methods to find reference transcriptomes with the highest similarity to query cluster expression profiles, peripheral blood mononuclear cell (PBMC) clusters are correctly labeled using either bulk-RNA seq references generated from the ImmGen database[9,16], processed microarray data of purified cell types[17], or previously annotated scRNA-seq results[11] (Fig. 1B). We reached similarly satisfactory results in scRNA-seq brain transcriptome data from mouse and human samples, as detailed by scRNAseq_Benchmark[14] (F1-score of 1 for all 4 identity mapping pairs, on 3 main cell types, data not shown).

To assess the performance of clustifyr, we used the Tabula Muris dataset[5], which contains data generated from 12 matching tissues using both 10x 3’ end seq (“drop”) and SmartSeq2 (“facs”) platforms. Using references built from “facs” Seurat objects, we attempted to assign cell type identities to clusters in “drop” Seurat objects. In benchmarking results, clustifyr is comparably accurate versus other automated classification packages (Fig. 1C). Cross-platform comparisons are inherently more difficult, and the approach used by clustifyr is aimed at being platform- and normalization-agnostic. Mean runtime, including both reference building and test data classification, in Tabular Muris classifications was ~ 1 second if the required variable gene list is extracted from the query Seurat object (Fig. 1D). Alternatively, variable genes can be recalculated by other methods such as M3Drop[15], to reach similar results.

We further benchmarked clustifyr against a suite of comparable datasets, PBMCbench[18], generated from 2 PBMC samples using multiple scRNA-seq methods. Notably, for each reference dataset cross-referenced to other samples, clustifyr achieved a median F1-score of above 0.94 using Spearman ranked correlation (Fig. 2A). Other correlation methods are on par or slightly worse at cross-platform classifications, which is expected based on the nature of ranked vs unranked methods. We therefore selected Spearman as the default method in clustifyr, with other methods also available, as well as a wrapper function to find consensus identities across available correlation methods (see Fig. 3B).

**FIGURE 2.**
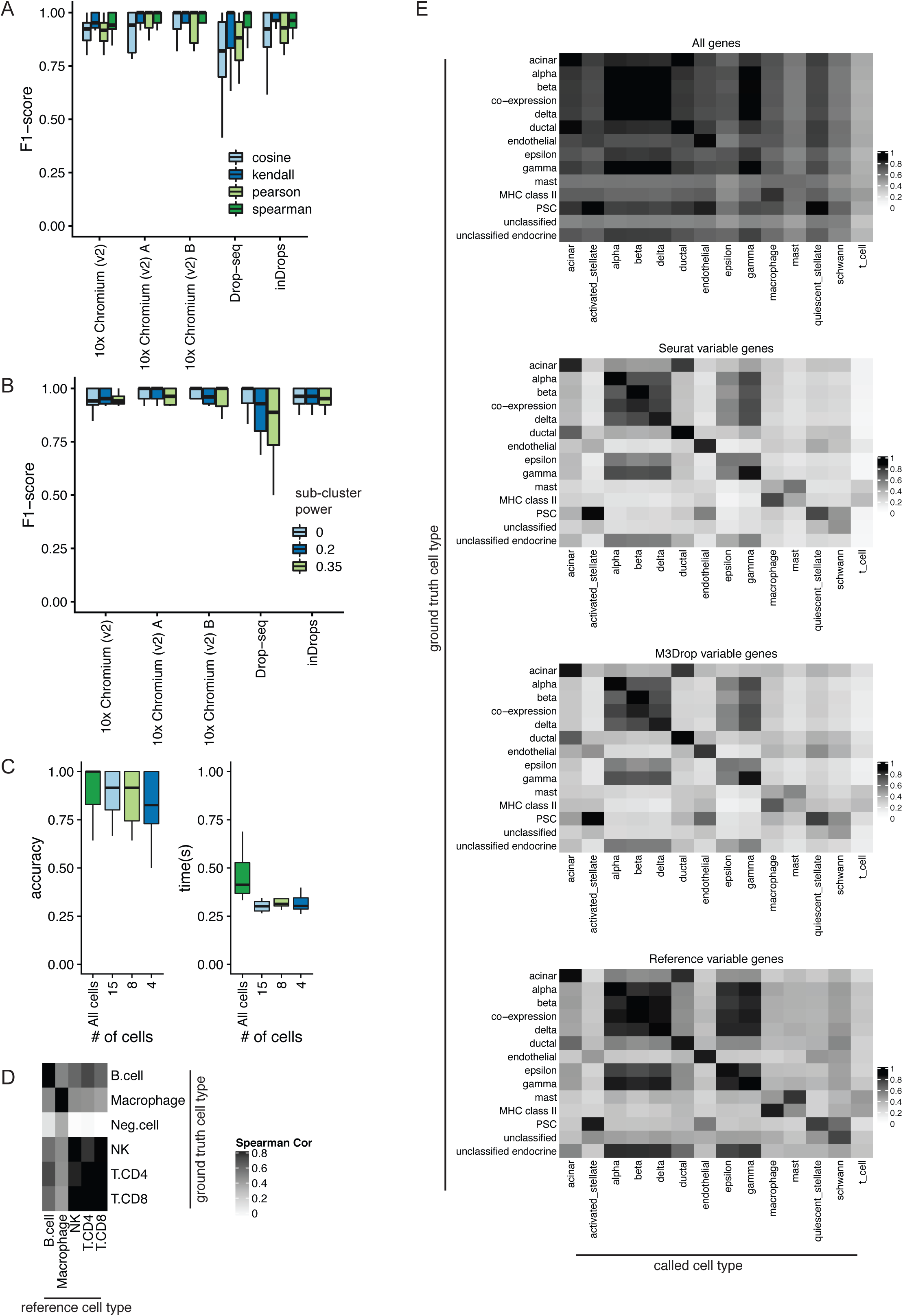
Parameter considerations for clustifyr. A) Comparison of accuracy of different correlation methods for classifying across platforms using the PBMCbench dataset. B) An assessment of the accuracy of using single or multiple averaged profiles as reference cell types was conducted using the PBMCbench test set. The number of reference expression profiles to generate for each cell type is determined by the number of cells in the cluster (n), and the sub-clustering power argument (x), with the formula n^x. C) Accuracy and performance were assessed with decreasing number of query cluster cell numbers using the PBMCbench test. D) Heatmap showing correlation coefficients between query cell types and the reference cell types. Clusters with correlation < 0.50 are assigned as Neg. Cell by clustifyr. E) Comparison of classification power using different feature selection methods (M3Drop, Seurat variable gene selection, selection of high variance genes from reference dataset, or no variable gene selection).

**FIGURE 3.**
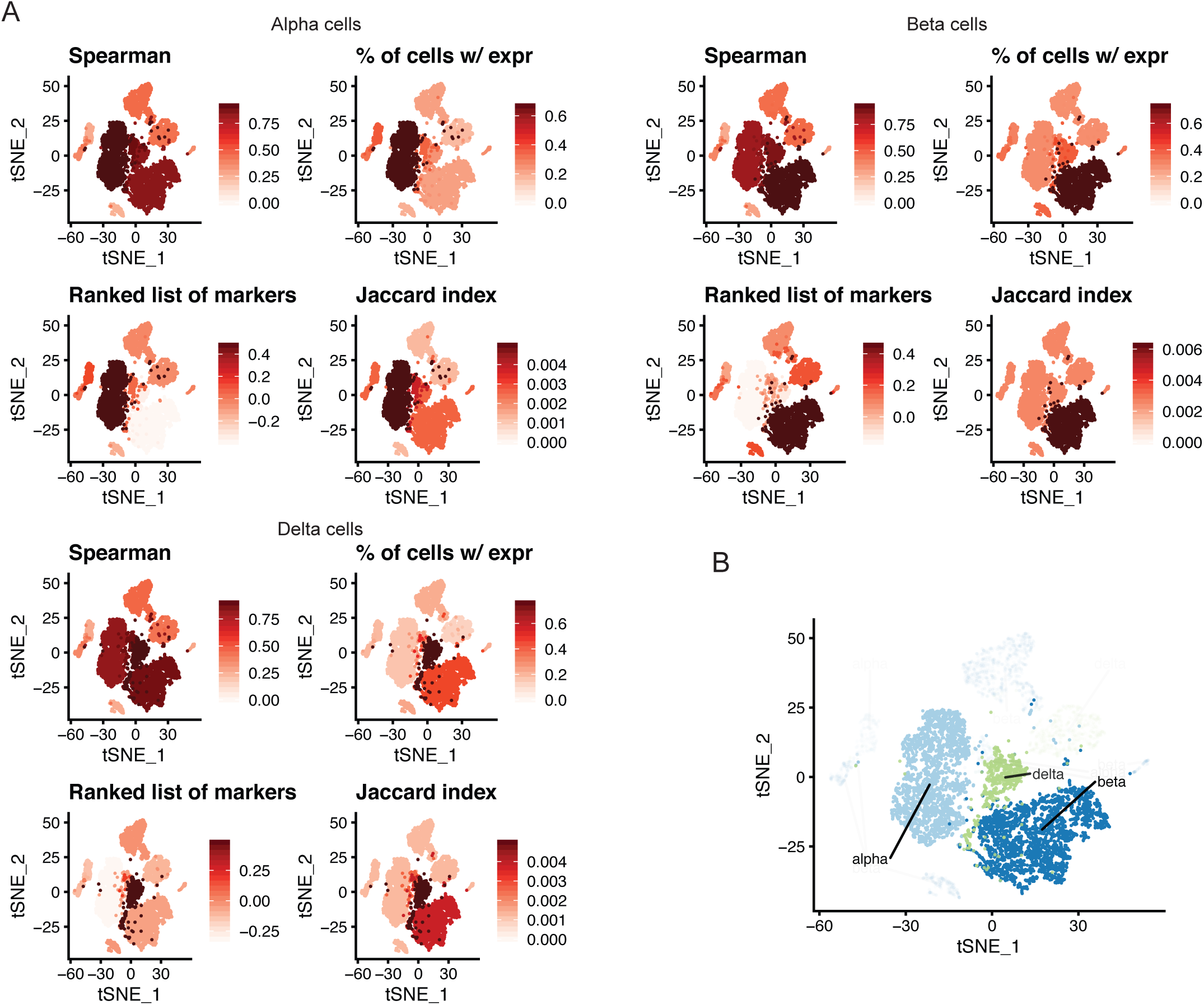
clustifyr implements multiple workflows for cell type classification. A) Comparison of ranked correlation vs gene list metrics for alpha, beta, and delta cells in pancreatic dataset. B) Consensus cell type calls from using a reference scRNA-seq dataset and gene list methods on alpha, beta, and delta cells in the pancreatic data.

For scRNA-seq reference data, matrices are built by averaging per-cell expression data for each cluster, to generate a transcriptomic snapshot similar to bulk RNA-seq or microarray data. An additional argument to subcluster the reference dataset clusters is also available, to generate more than one expression profile per reference cell type. The number of subclusters for each reference cell type is dependent on the number of cells in the cluster (n), and the sub-clustering power argument (x), following the formula n^x[9]. This approach does not improve classification in the PBMCbench data (Fig. 2B), however. We envision its utility would greatly depend on the granularity of the clustering in the reference dataset.

We also tested a general reference set built from the Mouse Cell Atlas[19], and found classification of the Tabula Muris data to be of high accuracy (Fig. 2C). Therefore, clustifyr is useful in identity-mapping across different techniques, or simple exploratory analysis using generalized pre-built references. As expected, with further downsampling of the number of cells in each query cluster, we observe decreased accuracy. Yet, even at 15 cells per tested cluster, clustifyr still performed well, with a further increase in speed. Based on these results, we set the default parameters in clustifyr to exclude classification of clusters containing less than 10 cells.

Recognition of missing reference cell types, so as to avoid misclassification, is another point of great interest in the field. From general usage of clustifyr, we find using a minimum correlation cutoff of 0.5 or 0.4 is generally satisfactory. Alternatively, the cutoff threshold can be determined heuristically using 0.8 * highest correlation coefficient among the clusters. One example is shown in Fig. 2D, using benchmark data modified by the SciBet package[20]. Megakaryocytes were removed from reference data, and labeled as “neg.cells” for ground truth in test data. clustifyr analysis found the “neg.cells” to be dissimilar to all available reference cell types, and hence left as “unassigned” under the default minimum threshold cutoff. Next, we applied clustifyr to a series of increasingly challenging datasets from the scRNAseq_Benchmark[14] unseen population rejection test. Without the corresponding cell type references, 57.5% of T cells were rejected and unassigned. When only CD4+ references were removed, 28.2% of test CD4+ T cells were rejected and unassigned. clustifyr was unable to reject CD4+/CD45RO+ memory T cells, mislabeling them as CD4+/CD25 T Reg instead when the exact reference was unavailable. However, these misclassifications are also observed with other classification tools benchmarked in the scRNAseq_Benchmark study[14].

As the core function of clustifyr is ranked correlation, feature selection to focus on highly variable genes is critical. In Fig. 2E, we compare correlation coefficients using all detected genes (>10,000), feature selection by the package M3Drop, variable genes selected by Seurat VST (default takes top 2,000), and using 1,000 genes with highest variance in the reference data. As seen, a basic level of feature selection is sufficient to classify the pancreatic cells. In the case of other cell type mixtures, especially ones without complete knowledge of the expected cell types, clustering and feature selection will be of greater importance. clustifyr does not provide novel clustering or feature selection methods on its own, but instead is built to maintain flexibility to incorporate methods from other, and future, packages. We view these questions as fast-moving fields[21,22], and hope to benefit from new advances, while keeping the general clustifyr framework intact.

Reliable and high-quality full transcriptome datasets are often not available for many cell types and therefore biologists must use a short list of marker genes established from literature to identify cell types. To replace the inefficient experience of plotting the expression of a handful of key marker genes and manually assigning cell types, clustifyr also implements quick methods of gene list enrichment analysis. Using ranked and unranked lists, respectively, clustifyr can correctly annotate PBMC and pancreas scRNA-seq clusters (data not shown). We tested the gene list functionality of clustifyr against the same test of 12 Tabula Muris reference and test pairs, as described above for the ranked correlation approach. With automated marker gene selection, ~85% of clusters were classified correctly (clustifyr_lists in Fig. 1C). In real world use cases, we expect the marker gene lists to be more carefully tailored, and hence perform better. In Fig. 3A, we compare the various calculated metrics of clustifyr, using ranked correlation on variable genes or a list of 5 previously established markers, and observe a consensus result identifying the alpha, beta, and delta cell clusters correctly. To combine all analysis, a function assesses consensus results across multiple classification methods within clustifyr and plots consensus cell types (Fig. 3B).

## CONCLUSIONS

We present a flexible and lightweight R package for cluster identity assignment. The tool bridges various forms of prior knowledge and scRNA-seq analysis. Reference sources can include scRNA-seq data with cell types assigned (or average expression per cell type, which can be stored at much smaller file sizes), sorted bulk RNA-seq, microarray data, and ranked or unranked gene lists. clustifyr, with minimal package dependencies, is compatible with a number of standard analysis workflows such as Seurat or Bioconductor, without requiring the user to perform the error-prone process of converting to a new scRNA-seq data structure, and can be easily extended to incorporate other data storage object types. Benchmarking reveals the package performs well in mapping cluster identity across different scRNA-seq platforms and experimental types.

On the user end, clustifyr is built with simple out-of-the-box wrapper functions, sensible defaults, yet also extensive options for more experienced users. Instead of building an additional single-cell-specific data structure, or requiring specific scRNA-seq pipeline packages, it simply handles basic data.frames (tables) and matrices (Fig. 1A). Input query data and reference data are intentionally kept in expression matrix form for maximum flexibility, ease-of-use, and ease-of-interpretation. Also, by operating on predefined clusters, clustifyr has high scalability and minimal resource requirements on large datasets. Using per-cluster expression averages results in rapid classification. However, cell-type annotation accuracy is therefore heavily reliant on appropriate selection of the number of clusters. Users are therefore encouraged to explore cell type annotations derived from multiple clustering settings. Additionally, assigning cell types using discrete clusters may not be appropriate for datasets with continuous cellular transitions such as developmental processes, which are more suited to trajectory inference analysis methods. As an alternative, clustifyr also supports per-cell annotation, however the runtime is greatly increased and the accuracy of the cell type classifications are decreased due to the sparsity of scRNA-seq datasets, and requires a consensus aggregation step across multiple cells to obtain reliable cell type annotations.

To further improve the user experience, clustifyr provides easy-to-extend implementations to identify and extract data from established scRNA-seq object formats, such as Seurat[11], SingleCellExperiment[12], URD[4], and CellDataSet (Monocle)[23]. Available in flexible wrapper functions, both reference building and new classification can be directly achieved through scRNA-seq objects at hand, without going through format conversions or manual extraction. The wrappers can also be expanded to other single cell RNA-seq object types, including the HDF5-backed loom objects, as well as other data types generated by CITE-seq and similar experiments[24]. Tutorials are documented online to help users integrate clustifyr into their workflows with these and other bioinformatics software.

## AVAILABILITY

clustifyr is submitted for review as a Bioconductor package and is licensed under the MIT license. Up-to-date source code, tutorials, and prebuilt references are available at https://github.com/rnabioco/clustifyr. Data used in examples and prebuilt references can also be found at https://github.com/rnabioco/clustifyrdata.

## ABBREVIATIONS

PBMC: peripheral blood mononuclear cell
scRNA-seq: single-cell RNA sequencing
SCE: SingleCellExperiment

## AUTHOR’S CONTRIBUTIONS

**Software** RF, AEG, RMS, CT, MD, YH, JRH, KAR; **Conceptualization** RF, AEG, KAR; **Writing – original draft** RF**; Writing – review & editing** RF, AEG, RMS, CT, MD, YH, JRH, KAR; **Supervision** JRH;

